# Sirtuin 1 is an endogenous NETosis inhibitor that becomes dysfunctional in diabetes

**DOI:** 10.1101/2025.08.05.668601

**Authors:** Liang De Wang, Rinkoo Dalan, Han Wei Hou, Siu Ling Wong

## Abstract

Neutrophils release their chromatin with toxic granular proteins as neutrophil extracellular traps (NETs) when activated. Diabetes exacerbates NET formation (NETosis), resulting in tissue damage and diabetic complications such as non-healing wounds. How diabetes predisposes neutrophils to NETosis remains unclear. Herein, we found that pharmacological inhibition or siRNA-knockdown of sirtuin 1 (SIRT1) increased NETosis in neutrophils of healthy humans and mice, unveiling SIRT1 as an endogenous suppressor of NETosis. In contrast, SIRT1 inhibition did not cause further increase in NETosis in neutrophils of diabetic patients and mice, indicative of SIRT1 dysfunction in disease state. Indeed, SIRT1 activity was significantly lower in neutrophils of diabetic individuals, accompanied by a concomitant increase in the activity of peptidylarginine deiminase 4 (PAD4), a key enzyme that mediates NETosis. PAD4 was co-detected with SIRT1 immunoprecipitated from HL-60-derived neutrophils cultured in basal glucose; such co-immunoprecipitation was absent in cells with high-glucose exposure, suggesting that hyperglycemia disrupts the SIRT1-PAD4 interaction. SIRT1 activators restored the SIRT1-PAD4 interaction and normalized the exacerbated NETosis and PAD4 activity in diabetes and hyperglycemia. This study reveals a novel regulatory role of SIRT1 on PAD4 activity. Revitalizing SIRT1 can be a new preventive or therapeutic strategy for combating NET-mediated inflammation in diabetes and beyond.

## Introduction

Neutrophils represent an indispensable arm of innate immunity, where they execute host defense through phagocytosis and degranulation of antimicrobials such as myeloperoxidase. Activated neutrophils can also form the highly pro-thrombotic and pro-inflammatory NETs via a series of well-orchestrated cellular events.[1] Initially discovered to be host protective in limiting pathogen dissemination,[2] the deleterious role of NETs was soon recognized in various diseases and conditions in sterile inflammation.[3–5] Unwarranted NET formation occurs in neutrophils of humans and mice with diabetes despite an overt absence of infection, leading to delayed wound recovery, non-healing ulcers, diabetic retinopathy, and diabetic nephropathy.[6–9] Their neutrophils also exhibit a four-fold increase in the expression of peptidylarginine deiminase 4 (PAD4),[7] a Ca^2+^- dependent enzyme critical for NETosis.[10–11]

PAD4 decondenses chromatin by reducing electrostatic interactions between histones and DNA through converting the positively charged arginine residues on histones to neutral citrulline. [11] Unlike other PAD isoforms, PAD4 expression is mainly restricted to granulocytes, and PAD4 is the only member that contains a nuclear localization signal.[10] The critical role of PAD4 in NET formation has been validated using PAD4-deficient mice[12–14] and CRISPR/Cas9-mediated knockdown of *PADI4* in HL-60-derived human neutrophils (dHL-60 cells).[1]

While diabetes primarily results from disorders in metabolism, it is a biological aging process that shares similarities with chronological aging in which NET-implicated organ dysfunction becomes prominent.[15] Diseases and conditions that involve neutrophils or NETs, such as thrombosis and atherosclerosis, are prevalent in both aged individuals and patients with diabetes. Similar to neutrophils of mice and humans with diabetes, aged neutrophils are also more predisposed to NETosis.[15–16] Aging induces profound epigenetic reprogramming in neutrophils, many of which leads to aberrant gene expression that augments inflammation.[17] Evolutionarily conserved mechanisms such as sirtuins serve to mitigate the detrimental impact of genotoxic insults.[18]

Sirtuins are nicotinamide adenine dinucleotide (NAD+)-dependent histone deacetylases and ADP-ribosyltransferases that are non-redundant in regulating the physiology related to nutrition, cellular stress, and aging.[18] Identified in *Saccharomyces cerevisiae*, Silent Information Regulator 2 (Sir2) is the first sirtuin member that was shown to prolong lifespan by suppressing genome instability.[19–20] Seven members of sirtuins (SIRT1-7) have been described in mammals;[21] amongst which, SIRT1, the best characterized member, is a nuclear deacetylase whose canonical substrates include histones, and other non-histone targets such as the tumor suppressor protein p53[22] and NF-κB transcription factor.[23] In macrophages, deletion of SIRT1 results in excessive levels of hyperacetylated RelA/p65 subunit of NF-κB, which promotes transcriptional activation of proinflammatory genes.[24] The impact of SIRT1 in neutrophils, in particular NETosis in metabolic disease, has not been explored.

While NET release depends on glucose availability,[25] how excessive glucose and diabetes exacerbates NETosis remains undefined. Since diabetes could promote biological aging, we sought to address whether and how SIRT1 regulates NETosis in health and diabetes. Herein, using blood neutrophils freshly isolated from humans and mice with diabetes, as well as dHL-60 cells with prolonged high-glucose exposure, we demonstrate that SIRT1 limits NETosis in healthy neutrophils via PAD4 inhibition, and this “endogenous NETosis inhibitor” becomes dysfunctional in diabetes and hyperglycemia, thus resulting in excessive NET formation. We identified a novel regulatory role of SIRT1 on PAD4 activity. Acute treatment using SIRT1 activators such as resveratrol and SRT2104, can markedly reduce NETosis and PAD4 activity in diabetic conditions.

## Results

### Pharmacological inhibition or siRNA knockdown of SIRT1 exacerbates NETosis in healthy neutrophils

As shown in the microscopy-based NETosis assay (Figure S1, Supporting Information), neutrophils isolated from fresh blood of humans and mice could be stimulated to produce NETs using ionomycin (Figure 1A) and lipopolysaccharides (Figure 1B), respectively.[7] Thirty-minute pre-incubation of Ex-527 (SIRT1 inhibitor) increased stimulant-induced NET production in both healthy human and mouse neutrophils (Figure 1A,B), suggesting that SIRT1 may serve as an endogenous suppressor of NETosis. Ex-527 had no effect on the NETosis *per se* if the cells were not stimulated, as the levels of NETs generated by neutrophils treated with Ex-527 alone were similar to those of the unstimulated control (Figure 1A,B).

**Figure 1.**
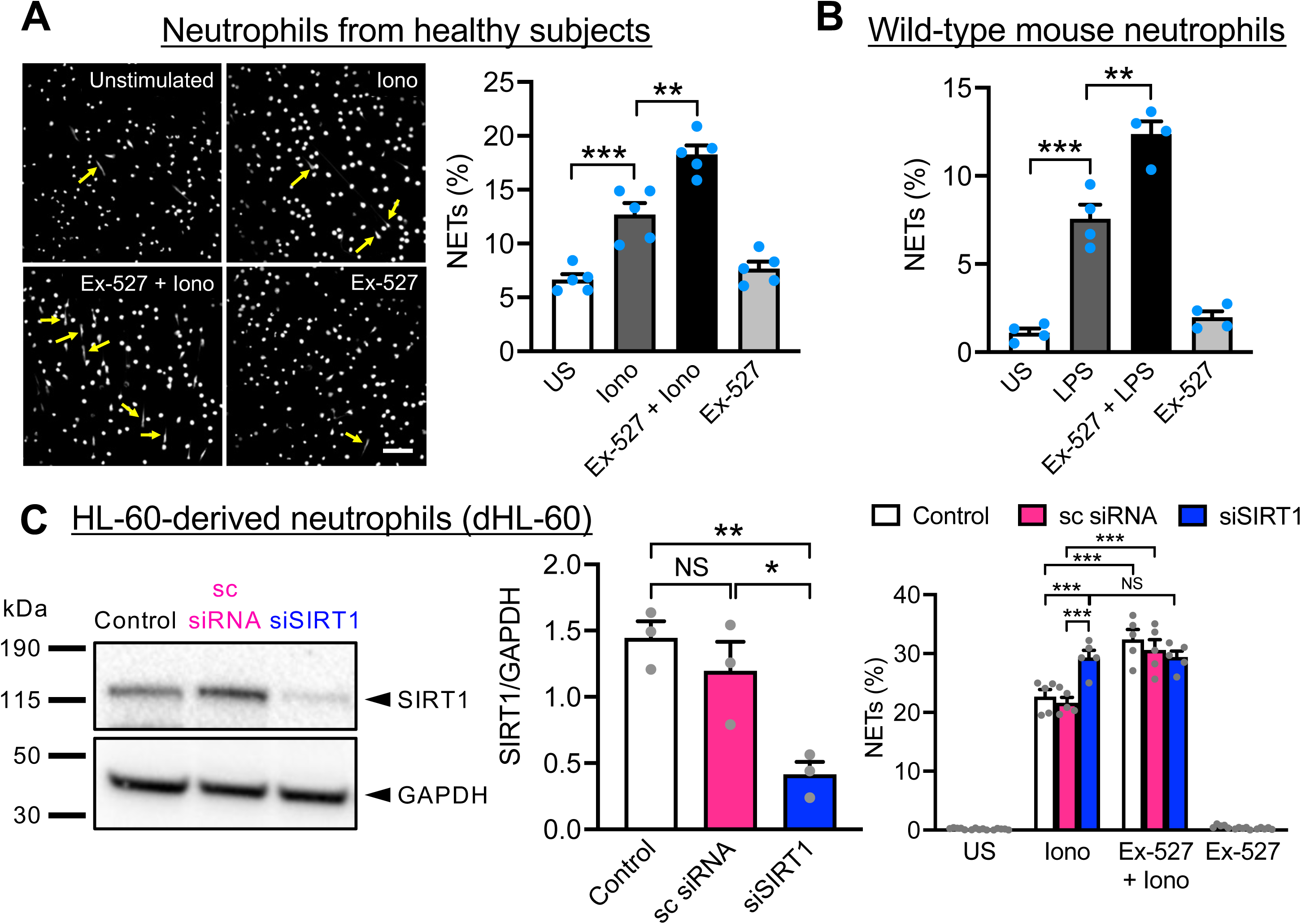
Pharmacological inhibition or siRNA knockdown of SIRT1 exacerbates NETosis in healthy neutrophils. (**A**) (Left) Representative images of ionomycin-induced NET formation in neutrophils isolated from healthy subjects and the effect of 30-minute pre-treatment of Ex-527 (specific SIRT1 inhibitor) on NETosis. Cells were fixed and stained with Hoechst 33342. NETs are indicated by yellow arrows. Scale, 100 μm. (Right) Quantitative data showing percentage of cells generated NETs under the various treatments. Data were analyzed by unpaired two-tailed Student’s t-test, n = 5 subjects, **P<0.01, ***P<0.001. (**B**) Quantitative data showing percentage of wild-type mouse neutrophils that produced NETs when stimulated with lipopolysaccharides (LPS) with and without pre-treatment of Ex-527. Data were analyzed by unpaired two-tailed Student’s t-test, n = 4 mice, **P<0.01, ***P<0.001. (**C**) (Left and middle) Western blot and summarized data showing levels of SIRT1 protein expression when HL-60-derived neutrophils (dHL-60 cells) were transfected with scrambled siRNA (sc siRNA) or siRNA targeting SIRT1 mRNA (siSIRT1). Data were analyzed by unpaired two-tailed Student’s t-test, n = 3 biological replicates, *P<0.05, **P<0.01, NS, non-significant. (Right) Effect of Ex-527 on dHL-60 cells with intact SIRT1 (non-manipulated control, and cells transfected with sc siRNA) and SIRT1- knockdown cells (siSIRT1). Data were analyzed by one-way ANOVA followed by Tukey’s multiple comparison, n = 5 biological replicates, ***P<0.001, NS, non-significant. US, unstimulated. Data are mean ± SEM.

To confirm the effect of Ex-527 was specific to SIRT1, we performed SIRT1 knockdown using siRNA on neutrophils differentiated from the human promyelocytic cell line HL-60 (dHL-60). Western blot analysis showed that SIRT1 expression was remarkably reduced in cells transfected with siRNA targeting SIRT1 (siSIRT1) (Figure 1C). Transfection with scrambled siRNA (sc siRNA) did not have an effect on SIRT1 expression, which was the same as the non-manipulated control (Figure 1C). When these cells were subjected to NETosis assay, similar to human and mouse neutrophils (Figure 1A,B), Ex-527 increased ionomycin-stimulated NETosis in both the non-manipulated control and dHL-60 cells that were transfected with sc siRNA (Figure 1C). Notably, excessive NETosis was already observed in dHL-60 cells transfected with siSIRT1 when they were only exposed to ionomycin, and addition of Ex-527 was without further effect (Figure 1C). The levels of NETosis in ionomycin-stimulated siSIRT1-transfected dHL-60 were comparable to those of control and sc siRNA-transfected cells treated with both ionomycin and Ex-527 (Figure 1C), strongly indicating that Ex-527 has no other targets except SIRT1, and SIRT1 is the key inhibitor in stimulant-activated NETosis.

### Exogenous SIRT1 inhibition is without effect in NETosis in diabetes

Since neutrophils of subjects and mice with diabetes are more susceptible to NETosis, we tested the hypothesis that exogenous SIRT1 inhibition could further increase NET formation in these hyperactivated neutrophils. We isolated blood neutrophils from individuals with diabetes and age-matched normoglycemic subjects. As previously reported,[6–7] NETosis levels were higher in neutrophils isolated from subjects with diabetes when compared to the normoglycemic controls (Figure 2A). Interestingly, while Ex-527 augmented ionomycin-stimulated NETosis in neutrophils isolated from normoglycemic controls, there was no further increase in NETosis in neutrophils isolated from subjects with diabetes (Figure 2A). In fact, diabetes *per se* mimics the effect of Ex- 527 treatment on NET formation. The NETosis levels of neutrophils from subjects with diabetes only singly stimulated with ionomycin were similar to that of the Ex-527-treated ionomycin-stimulated neutrophils of normoglycemic controls (Figure 2A). This may suggest that neutrophil SIRT1 is dysfunctional in diabetes, thus resulting in full-scale NETosis which is not aggravated by exogenous SIRT1 inhibition from Ex-527.

**Figure 2.**
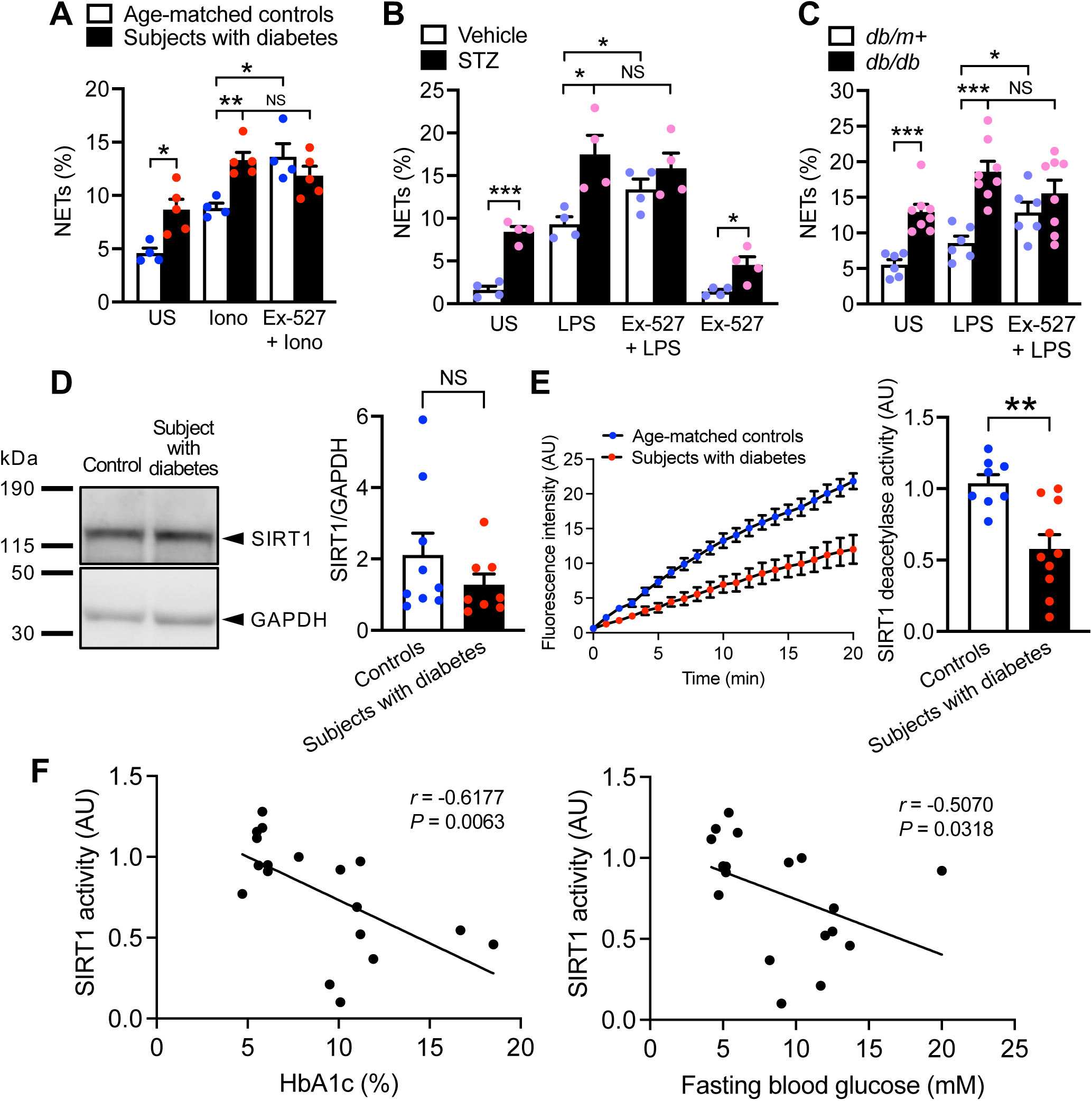
Exogenous SIRT1 inhibition has no effect on NETosis in neutrophils isolated from diabetic humans and mice. (**A-C**) Quantitative data showing percentage of NETting cells in neutrophils isolated from (**A**) subjects with diabetes and the age-matched healthy controls, (**B**) the streptozotocin (STZ)-induced type 1 diabetic mouse model, and (**C**) the *db/db* type 2 diabetic mice (and its normoglycemic *db/m+* control) when stimulated for NETosis with and without pre- treatment of Ex-527. Data were analyzed by unpaired two-tailed Student’s t-test, (**A**) n = 4 individuals for age-matched controls, n = 5 individuals for subjects with diabetes, (**B**) n = 4 mice per group, (**C**) n = 6 mice for *db/m+*, n = 8 mice for *db/db*, *P<0.05, **P<0.01, ***P<0.001, NS, non-significant. (**D**) (Left) Western blot and (right) summarized data showing levels of SIRT1 protein expression in neutrophils isolated from individuals with diabetes and the age-matched healthy subjects. Data were analyzed by unpaired two-tailed Student’s t-test, n = 5 individuals for age-matched controls, n = 4 individuals for subjects with diabetes. (**E**) (Left) Time-course plot and (right) the computed deacetylase activity of SIRT1 immunoprecipitated from equal number of neutrophils from subjects with diabetes and age-matched healthy individuals. Data were analyzed by unpaired two-tailed Student’s t-test, n = 8 individuals for age-matched controls, n = 10 individuals for subjects with diabetes, **P<0.01. (**F**) Two-tailed Spearman correlation analysis conducted between (left) HbA1c and SIRT1 activity and (right) fasting blood glucose and SIRT1 activity of blood neutrophils isolated from age-matched controls (n = 9) and subjects with diabetes (n = 10).

We also investigated the effect of Ex-527 in the neutrophils of STZ-induced diabetic mice and *db/db* mice, type 1 and type 2 diabetic models, respectively. Similar to human neutrophils (Figure 2A), Ex-527 increased LPS-stimulated NETosis in the vehicle-treated (Figure 2B) and *db/m+* (Figure 2C) normoglycemic controls, but not the STZ-induced diabetic mice nor the *db/db* mice (Figure 2B,C). The findings showed that the nullifying effect of diabetes on the “endogenous SIRT1 brake of NETosis” is universal and conserved across species.

SIRT1 expression and activity were further examined in the human neutrophils. Western blot analysis revealed that neutrophil SIRT1 expression was not significantly different between the normoglycemic subjects and individuals with diabetes (Figure 2D). However, when the SIRT1 protein was immunoprecipitated from equal number of neutrophils and examined for activity, neutrophils from individuals with diabetes exhibited remarkably lower SIRT1 activity (Figure 2E). In fact, correlation analysis revealed a significant negative relationship between blood glucose and SIRT1 activity (Figure 2F). The higher the blood glucose level, the lower was the SIRT1 activity (Figure 2F). SIRT1 activity exhibited a stronger correlation with HbA1c than with fasting blood glucose (Figure 2F), possibly because HbA1c was more accurate in reflecting the subjects’ hyperglycemic burden across time. The diminished endogenous suppression of NETosis resulting from a reduction of SIRT1 activity explains the exacerbated NETosis in diabetes.

### Hyperglycemia *per se* is sufficient to make SIRT1 defective

Since hyperglycemia is the common denominator of type 1 and type 2 diabetes, we sought to understand if high glucose contributed to SIRT1 dysfunction. Blood neutrophils are too short-lived *ex vivo* to examine the chronic impact of hyperglycemia as in diabetes, we thus optimized the dHL-60 cellular model for hyperglycemia-related investigations. After 5 days of differentiation towards the neutrophil lineage using 1.3% DMSO, dHL-60 cells either continued to be cultured in basal glucose (11 mM, standard medium for dHL-60 cell line) or switched to high glucose (44 mM) medium for 24 hours (Figure 3A). The cells were then seeded in DMSO-free basal glucose medium for NETosis assay. We found that high glucose-treated neutrophils had remarkably higher NETosis when compared to those cultured in basal glucose (Figure S2, Supporting Information). Mannitol supplementation instead of glucose (33 mM mannitol) was without effect (Figure S2, Supporting Information), indicating that osmotic pressure is not a cause of the change in NETosis under high glucose. The effect of high glucose was also specific to dHL-60 cells (i.e., neutrophils), but not to the native (non-differentiated) promyelocytes (Figure S2, Supporting Information), validating the dHL-60 cell line model for studying NETosis in hyperglycemic condition.

**Figure 3.**
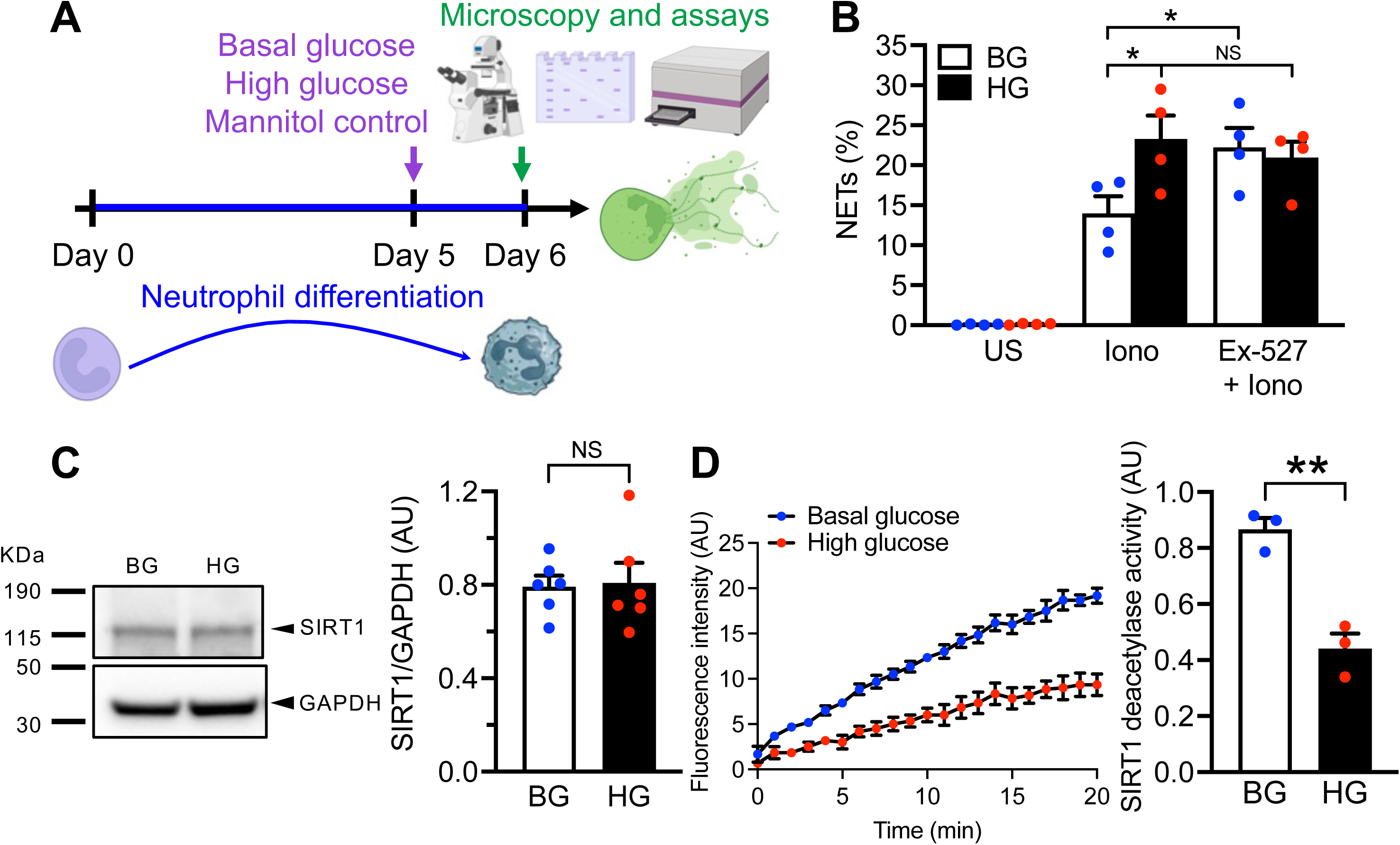
Hyperglycemia *per se* is sufficient in reducing neutrophil SIRT1 activity. (**A**) Schematic of experimental design (created with BioRender.com) for examining the impact of high glucose on the NETosis inhibitory effect of SIRT1. (**B**) Quantitative data showing percentage of basal glucose (BG, 11 mM)- or high glucose (HG, 44 mM)-cultured dHL-60 cells that produced NETs when stimulated for NETosis with and without pre-treatment of Ex-527. Data were analyzed by unpaired two-tailed Student’s t-test, n = 4 biological replicates per group, *P<0.05, NS, non-significant. (**C**) (Left) Western blot and (right) summarized data (right) showing levels of SIRT1 protein expression in BG- or HG-cultured dHL-60 cells. Data were analyzed by unpaired two-tailed Student’s t-test, n = 6 biological replicates per group, NS, non-significant. (**D**) (Left) Time-course plot and (right) the computed deacetylase activity of SIRT1 immunoprecipitated from equal number of BG- or HG-cultured dHL-60 cells. Data were analyzed by unpaired two-tailed Student’s t-test, n = 3 biological replicates per group, **P<0.01. Data are mean ± SEM.

Similar to blood neutrophils, Ex-527 only increased stimulant-activated NETosis in cells cultured in basal glucose, but not the high glucose-exposed cells (Figure 3B). Twenty-four hours of high glucose exposure did not alter SIRT1 expression (Figure 3C), but markedly reduced SIRT1 activity (Figure 3D). These observations implicate that high glucose is a negative modulator of SIRT1 activity and impairs NETosis-inhibitory function of SIRT1 in diabetes.

### SIRT1 dampens NETosis by suppressing PAD4 activity

PAD4 is a critical enzyme for NETosis.[1, 12–14] We therefore investigated whether there is any relationship between SIRT1 and PAD4. Activity of PAD4 in the neutrophils was evaluated by immunostaining of citrullinated histone H4 (H4Cit). When stimulated, Ex-527 increased the number of H4Cit^high^ neutrophils isolated from normoglycemic subjects (Figure 4A) and mice (Figure 4B–D), as well as dHL-60 cultured in basal glucose (Figure 4E). However, such increase was not observed in neutrophils in diabetes (Figure 4A–D) nor hyperglycemia (Figure 4E). These strongly suggest that PAD4 is downstream of SIRT1, which reduces NETosis by suppressing PAD4 activity when it is functional in health. While a strong significant positive correlation was observed between HbA1c and levels of H4Cit^high^ cells (Figure 4F), SIRT1 activity negatively correlated with levels of H4Cit^high^ cells (Figure 4F), aligning with our findings that diabetes and the reduction of SIRT1 activity both increase PAD4 activity (Figure 4A). To confirm the NETosis inhibitory effect of SIRT1 is indeed mediated through PAD4 inhibition, we performed NETosis assay on blood neutrophils isolated from mice that were deficient in PAD4 in their hematopoietic cells (*Vav1-Cre Padi4^fl/fl^*). As expected, Ex-527 elevated NETosis in neutrophils from the control mice (*Padi4^fl/fl^*), but such elevation was not observed in the PAD4-deficient neutrophils (Figure 4G). Taken together, SIRT1 inhibits NETosis via suppression of PAD4 activity in both healthy human and mouse neutrophils.

**Figure 4.**
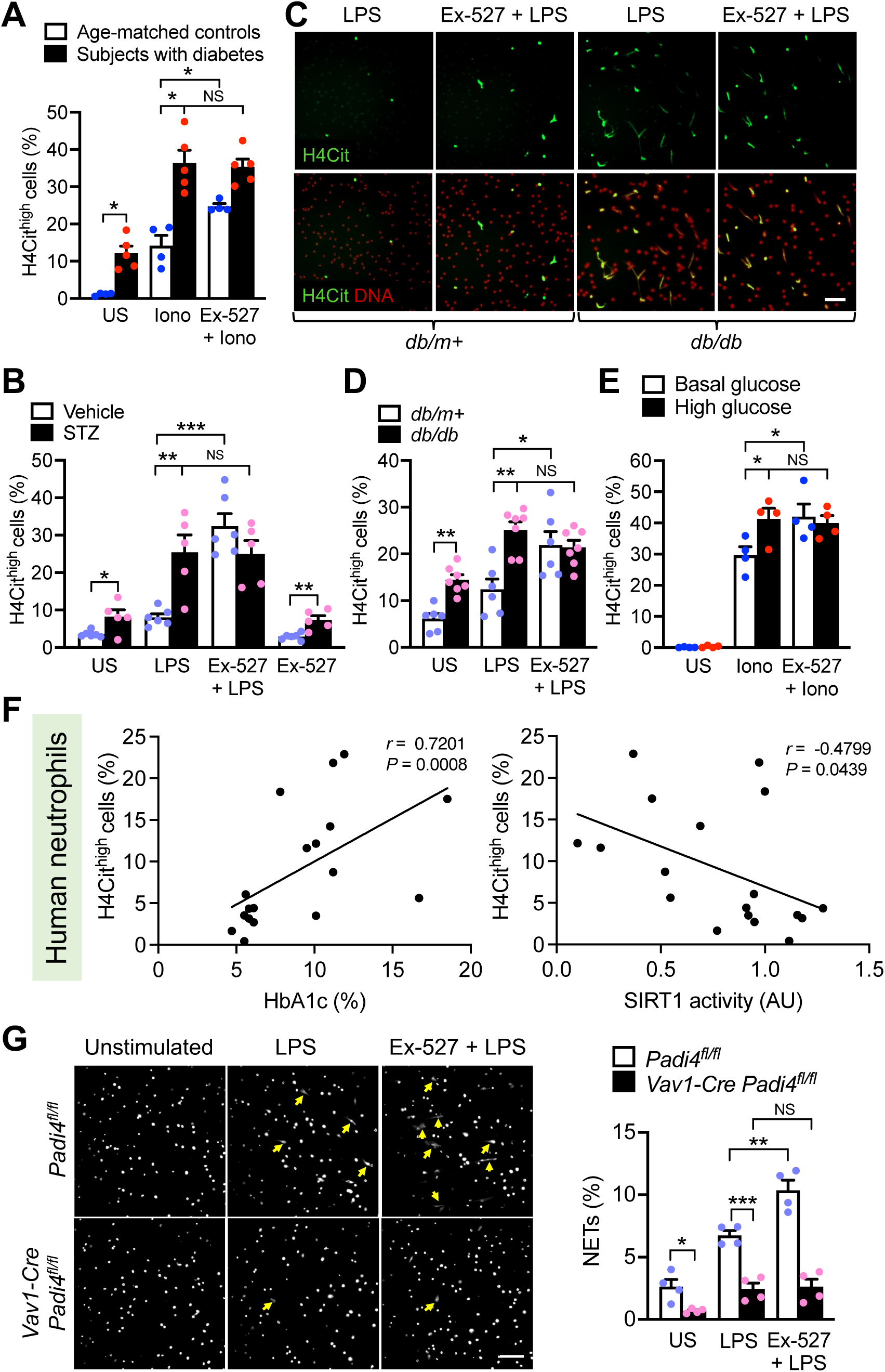
Ex-527 increases PAD4 activity in healthy neutrophils but not those of humans and mice with diabetes. (**A-E**) Histone H4 citrullination (H4Cit) was used as a marker for PAD4 activity. Percentage of H4Cit^high^ cells were quantified after neutrophils isolated from (**A**) subjects with diabetes and the age-matched healthy controls, (**B**) STZ-induced diabetic mice (and their normoglycemic vehicle-treated control), and (**C,D**) the diabetic *db/db* mice (and its normoglycemic *db/m+* control) were stimulated for NETosis with and without pre-treatment of Ex-527. (**C**) Representative images showing the effect of Ex-527 on H4Cit in LPS-stimulated neutrophils of *db/m+* and *db/db* mice. Scale, 100 μm. (**E**) Percentage of H4Cit^high^ cells were quantified in basal and high glucose-treated dHL-60 cells when stimulated for NETosis in the presence or absence of Ex-527. Data were analyzed by (**A**) Mann-Whitney test, n = 4 individuals for age-matched control, n = 5 individuals for subjects with diabetes, (**B**) unpaired two-tailed Student’s t-test, n = 6 mice for vehicle, n = 5 mice for STZ, (**D**) Mann-Whitney test, n = 6 mice for *db/m+*, n = 7 mice for *db/db*, and (**E**) unpaired two-tailed Student’s t-test, n = 4 biological replicates per group. (**F**) Two-tailed Spearman correlation analysis performed between (left) HbA1c and levels of H4Cit^high^ cells and (right) SIRT1 activity and levels of H4Cit^high^ cells of blood neutrophils isolated from age- matched controls (n = 9) and subjects with diabetes (n = 10). (**G**) (Left) Representative images of NET formation in neutrophils isolated from hematopoietic cell-specific PAD4-deficient mice (*Vav1- Cre Padi4^fl/fl^*) and the control (*Padi4^fl/fl^*) in response to Ex-527 during NETosis. NETs are indicated by yellow arrows. Scale, 100 μm. (Right) Quantitative data showing percentage of neutrophils released NETs. Data were analyzed by unpaired two-tailed Student’s t-test, n = 4 mice per group (both male and female mice were used in each group). *P<0.05, **P<0.01, ***P<0.001, NS, non- significant. Data are mean ± SEM.

### SIRT1 activators, resveratrol and SRT2104, reduce NETosis in diabetes and hyperglycemia

To examine whether SIRT1 (re)activation can lessen the aberrant NETosis in diabetes, blood neutrophils isolated from humans and mice with diabetes were pre-incubated for an hour with resveratrol (RSV, 50 μM), a natural non-flavonoid polyphenol which is known to activate SIRT1 by binding to the allosteric site,[26–27] or SRT2104 (3 μM), a second generation of specific, small molecule SIRT1 activator that has been advanced to clinical trials,[28] before stimulated for NETosis. Remarkably, RSV and SRT2104 prevented stimulant-induced NETosis in neutrophils from humans and mice with diabetes, as well as dHL-60 cells treated with high glucose (Figure 5A–C,G). The effect of RSV and SRT2104 is likely mediated via PAD4 inhibition, as reflected by a concomitant reduction in H4Cit^high^ neutrophils after the treatments (Figure 5D–F,H). Specificity of RSV and SRT2104 was confirmed using SIRT1-knockdown dHL-60 cells, which demonstrated that both compounds had no effects on NETosis and histone citrullination (PAD4 activity) in the absence of SIRT1 (Figure 5G,H). In contrast to RSV and SRT2104, co-substrate supplementation with β-nicotinamide adenine dinucleotide (β-NAD, 100 μM, 30-minute pre-incubation) was without effect (Figure S3, Supporting Information).

**Figure 5.**
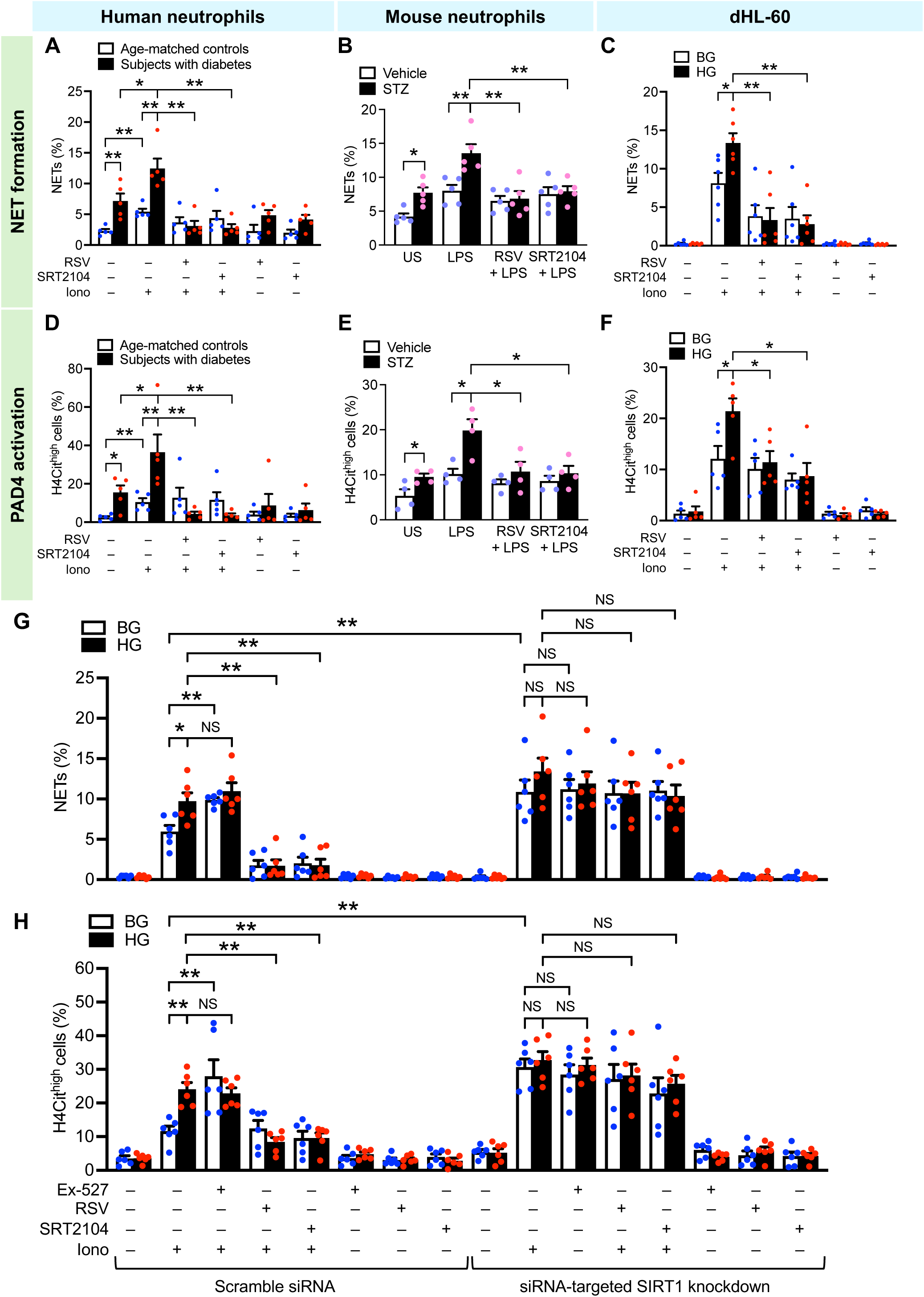
SIRT1 activators, resveratrol (RSV) and SRT2104, inhibit NETosis and histone hypercitrullination to comparable extent. (**A-C,G**) NETting cells and (**D-F,H**) H4Cit^high^ cells were quantified in neutrophils isolated from (**A,D**) subjects with diabetes (and the age-matched healthy controls), and (**B,E**) STZ-induced diabetic mice (and their normoglycemic vehicle-treated control), as well as high glucose-cultured dHL-60 cells (and the basal glucose-cultured counterparts) (**G,H**) with and (**C,F**) without SIRT1 knockdown procedures. Cells were pre-treated with Ex-527 (10 μM, 30 minutes), RSV (50 μM, 1 hour) or SRT2104 (3 μM, 1 hour) before stimulation with ionomycin or LPS, where indicated. Data were analyzed by (**A,D**) Mann-Whitney test, n = 5 individuals for age-matched controls, n = 5 individuals for subjects with diabetes, *P<0.05, **P<0.01; (**B,E**) Mann-Whitney test and unpaired two-tailed Student’s t-test, respectively, n = 4-5 mice for vehicle, n = 4-5 mice for STZ, *P<0.05, **P<0.01; (**C,F**) Mann-Whitney test, n = 5-6 biological replicates per group, *P<0.05, **P<0.01; (**G,H**) Mann-Whitney test, n = 6 biological replicates per group, *P<0.05, **P<0.01, NS, non-significant. Data are mean ± SEM.

### SIRT1-PAD4 interaction is disrupted by hyperglycemia

The SIRT1-specific effects of Ex-527, RSV and SRT2104 on histone citrullination (Figure 4, 5D– F,H) suggested that SIRT1 activity impacted on PAD4 activity, the latter being crucial for chromatin decondensation prior to NETosis.[1] Given that SIRT1 can also modulate chromatin conformation via histone deacetylation, we examined whether the inhibitory effect of SIRT1 on PAD4 was via SIRT1’s action on chromatin compaction *per se* (hence leading to reduced chromatin access by PAD4) or whether it was a direct interaction with PAD4. We examined acetylation of histone H3 in dHL-60 cells and found that levels of acetyl-H3^high^ cells were similar between cells cultured in basal glucose and those exposed to high glucose (Figure 6A), suggesting that the loss of SIRT1 activity under hyperglycemic condition (Figure 3D) does not directly open up the chromatin due to the loss of histone deacetylase activity. We then sought to examine if SIRT1 directly interact with PAD4 to inhibit PAD4 activity. High resolution confocal microscopy showed that both SIRT1 and PAD4 localized in the nucleus of dHL-60 cells (Figure 6B). Interaction between SIRT1 and PAD4 was confirmed by Western blot analysis. When SIRT1 was pulled down from dHL-60 cells, PAD4 was co-detected with the immunoprecipitated SIRT1 in cells cultured in basal glucose (Figure 6C). While Western blot of whole cell lysates for PAD4 and SIRT1 showed that both proteins were similarly expressed in basal glucose-cultured and high glucose-exposed cells (Figure 6C), PAD4 was not co-detected with the SIRT1 protein immunoprecipitated from dHL-60 cells exposed to high glucose (Figure 6C), indicating that the SIRT1-PAD4 interaction was disrupted by hyperglycemia. Intriguingly, SRT2104 restored the SIRT1-PAD4 interaction in high glucose-exposed cells (Figure 6D), suggesting that SIRT1 activity is crucial for SIRT1-PAD4 interaction.

**Figure 6.**
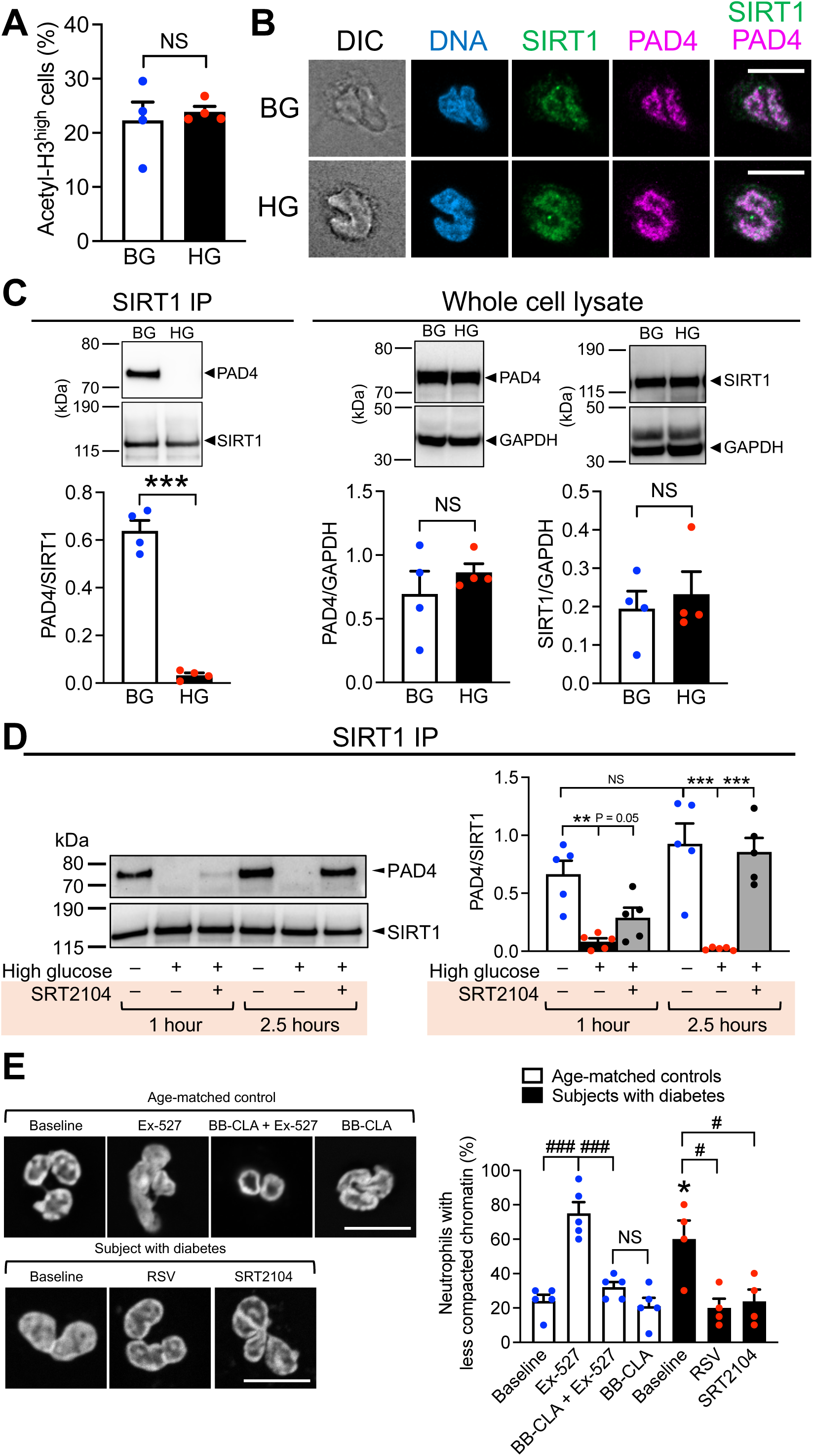
SIRT1-PAD4 interaction is disrupted by hyperglycemia. (**A**) Quantification of acetyl- H3^high^ dHL-60 cells with and without high glucose exposure using immunofluorescence microscopy. Data were analyzed by unpaired two-tailed Student’s t-test, n = 4 biological replicates per group, NS, non-significant. (**B**) Immunostaining of SIRT1 and PAD4 in dHL-60 cells cultured in basal glucose or primed in high glucose. Scale, 10 μm. (**C**) Western blot analysis of PAD4 and SIRT1 in (left) SIRT1 immunoprecipitated from dHL-60 cells and (right) whole cell lysate of dHL- 60. Data were analyzed by unpaired two-tailed Student’s t-test, n = 4 biological replicates per group. ***P<0.001, NS, non-significant. (**D**) dHL-60 cells were cultured in basal glucose or high glucose for 24 hours. High glucose-exposed cells were then treated with SRT2104 for 1 or 2.5 hours. (Left) Representative Western blot and (right) quantification showing PAD4 and SIRT1 levels in the SIRT1-immunoprecipitated samples from the dHL-60 cells. Data were analyzed by unpaired two-tailed Student’s t-test, n = 5 biological replicates per group. **P < 0.01, ***P<0.001, NS, non-significant. (**E**) Super-resolution imaging on chromatin compaction states in neutrophils isolated from age-matched healthy controls and subjects with diabetes. Cells were pre-treated with Ex-527 (10 μM, 30 minutes), RSV (50 μM, 1 hour), SRT2104 (3 μM, 1 hour), or BB-Cl-amidine (BB-CLA, 30 min), where indicated. BB-CLA was added 30 minutes prior to Ex-527 exposure when used in combination. Data were analyzed by unpaired two-tailed Student’s t-test, n = 5 individuals for age-matched controls, n = 4 individuals for subjects with diabetes, *P<0.05 compared to baseline of age-matched controls, ^#^P<0.05, ^###^P<0.001, NS, non-significant, compared between groups indicated. Data are mean ± SEM.

Since our data suggested that SIRT1 is constitutively active in neutrophils and dHL-60 cells and its activity is impaired by diabetes or high glucose treatment as reflected by the SIRT1 activity assay (Figure 2E,3D), we adopted super-resolution imaging to examine chromatin compaction states (Figure S4, Supporting Information) in various conditions where SIRT1 activity was altered. We isolated blood neutrophils from healthy individuals and subjects with diabetes and found that chromatin conformation was less compact in neutrophils isolated from subjects with diabetes, when compared to those from the age-matched healthy controls (Figure 6E). Ex-527 treatment *ex vivo* relaxed the compact heterochromatin in neutrophils of the healthy subjects, while RSV and SRT2104 restored chromatin compaction in neutrophils of subjects with diabetes (Figure 6E). Notably, when neutrophils from healthy subjects were pre-treated with BB-Cl-amidine (a PAD4 inhibitor) before exposing to Ex-527, the chromatin decondensing effect of Ex-527 was abrogated (Figure 6E). This series of *ex vivo* experiments collectively suggested that PAD4 is constitutively suppressed by direct inhibition from SIRT1 in healthy neutrophils.

## Discussion

Overall, we unveiled that SIRT1 is a natural, endogenous inhibitor of NETosis in health, and it becomes dysfunctional in diabetes. We identified hyperglycemia as a posttranslational modulator of SIRT1, which dampens NETosis via direct suppression of PAD4 activity. Notably, the “endogenous NETosis inhibitor”, SIRT1, can be revitalized by acute treatment of RSV and SRT2104 (Figure 7).

**Figure 7.**
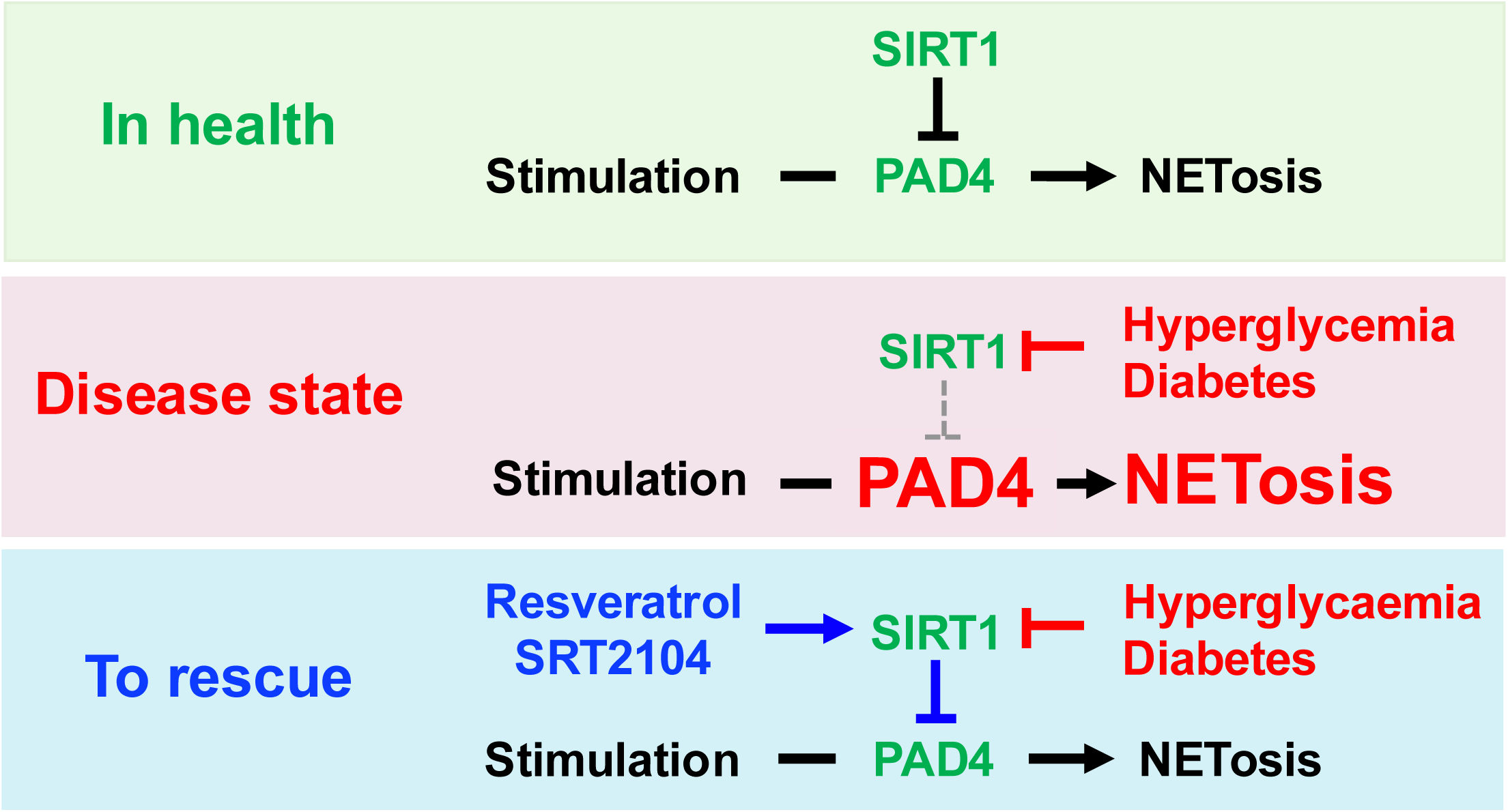
Schematic summary of findings of the study. SIRT1 serves as a natural, endogenous suppressor of NETosis in health via PAD4 inhibition. Diabetes and prolonged exposure to high glucose reduce SIRT1 activity, leading to an increase in PAD4 activity and excessive NET generation. To rescue, resveratrol and SRT2104 can acutely boost SIRT1 activity, thus lessening PAD4 activity and normalizing over-production of NETs in diabetes and hyperglycemia.

Changes in chromatin conformation is a crucial step prior to NET release. While neutrophil elastase and myeloperoxidase are implicated in chromatin decondensation,[29] PAD4 is a major driver in the massive unraveling of chromatin, particularly in the context of sterile inflammation.[5] Processes that counteract chromatin decondensation would make lytic NETosis more difficult. One of the major classes of enzymes that keep the chromatin in compacted state are sirtuins. SIRT1 is one of the class III histone deacetylases that modulate chromatin structure and thereby gene transcription. Shown both *in vitro* and *in vivo*, SIRT1 preferentially deacetylases histones H3 (H3K9 and H3K56) and H4 (H4K6), leading to chromatin silencing and repression of gene transcription.[30–31] However, how SIRT1 affects chromatin decondensation and therefore NETosis has not been explored. We presented the first evidence using PAD4-deficient mouse neutrophils that the inhibitory role of SIRT1 on NETosis is via inhibition of PAD4. Unlike wild-type blood neutrophils, those isolated from *Vav1-Cre Padi4^fl/fl^* mice failed to increase NETosis when treated with Ex-527, the SIRT1 inhibitor. In drastic contrast to SIRT1, deficiency of SIRT3, which localizes to the mitochondria and is important for downregulating mitochondrial reactive oxygen species by activating superoxide dismutase 2,[32–33] has no impact on NETosis despite a slight increase in intracellular oxidative stress [34]. All these data support that in the context of NETosis, the role of SIRT1 likely lies in chromatin structure modification, rather than its pleiotropic anti-oxidant property. Importantly, while SIRT1 can modulate chromatin compaction states by its histone deacetylase activity, our study showed that SIRT1’s impact on chromatin conformation is effected via PAD4. We revealed that SIRT1 inhibition (by Ex-527) did not result in chromatin decondensation when PAD4 was also inhibited (by BB-Cl-amidine). Co-detection of PAD4 with immunoprecipitated SIRT1 in basal glucose-cultured cells but not those treated in high glucose suggested that high glucose disrupts SIRT1-PAD4 interaction, liberating PAD4 to unravel chromatin without SIRT1’s suppression on its activity. This speculation is further supported by the findings that reactivation of SIRT1 activity by SRT2104 restores the SIRT1-PAD4 interaction, which results in a reduction in H4 hypercitrullination and hence NETosis. How high glucose intercept the SIRT1-PAD4 interaction, and how SIRT1 post-translationally modify PAD4 requires separate studies involving extensive proteomic analysis on acetylation sites of PAD4, site-directed mutagenesis, and functional confirmation. Since histone citrullination is one of the earlier events in NET formation [1] and we defined that SIRT1 is upstream of PAD4, the present findings could provide a handle for neutrophil programming by “rejuvenating” neutrophils via SIRT1 activation.

Metabolic disease has been reported to modulate SIRT1. SIRT1 mRNA and protein expression were reduced in peripheral blood mononuclear cells of humans with insulin resistance and metabolic syndrome.[35] High glucose and saturated fatty acids lower SIRT1 expression and diminish NAD+ levels in human monocytes.[35] In white adipose tissue, high fat diet can activate NLRP3 inflammasome-dependent caspase 1-mediated proteolytic cleavage of SIRT1.[36] Reduction in adipose SIRT1 expression is associated with an increase in adipose tissue inflammation in obese rodents and humans.[37] Chronic exposure of human vascular endothelial cells to high glucose *in vitro* reduces mRNA expression of all *SIRT* genes (*SIRT1-7*).[38–39] SIRT1 expression is also downregulated in retinal and renal tissue of diabetic mice and humans.[38, 40]

Intriguingly, we found that SIRT1 expression of blood neutrophils of individuals with diabetes and high-glucose exposed HL-60-derived neutrophils were unchanged, as opposed to other cell types. Diabetes and high glucose, however, markedly reduces SIRT1 activity. The reduction in neutrophil SIRT1 activity is unlikely due to depletion of NAD+, as our data showed that β-NAD supplementation does not restore SIRT1’s inhibition to NETosis in neutrophils of diabetic mice. How diabetes and high glucose inhibit SIRT1 activity posttranslationally requires further study. Ways to prevent or reverse posttranslational modification of SIRT1 could be instrumental for developing novel anti-NETosis therapeutics.

In this study, we also showed that RSV and SRT2104 (a natural and a more specific synthetic SIRT1 activator, respectively) could reduce exacerbated NETosis in diabetes. There is good potential to develop RSV and SRT2104 into therapeutic solutions for NET-mediated diabetic complications. However, translating our findings by taking the SIRT1 activators as a nutraceutical, as it has been, would be challenging and ineffective. Most of the RSV dose, for example, will be deactivated in the liver and the intestine when administered orally, resulting in low bioavailability at target site. In fact, our *ex vivo* data showing that NETosis can be decreased by acute treatment (one-hour pre-exposure) of RSV and SRT2104 supports the idea of turning the nutraceutical into a drug candidate. One should target the delivery of RSV or SRT2104 to sites of inflammation, particularly tissues with profound neutrophil infiltration, for optimal suppression of PAD4 and therefore NETosis. Preventing NETosis, and cleavage of NETs, are possible therapeutic approaches to curtail NET-mediated ailments. Nonetheless, clearance of the DNA backbone of NETs (for example, by DNase 1) is ineffective in removing other cytotoxic NET components including histones and neutrophil elastase,[41] which would remain in the tissue and continue propagating inflammation. Recently, small fragments of NETs have been reported to be immunogenic,[42] indicating that NET degradation may not be the best approach. Thus, ideally, anti-NET treatments should focus on preventing NET generation in the first place. Nonetheless, there is currently no NETosis inhibitor that has a good toxicity profile for humans.[43] Our novel findings of SIRT1 being an effective suppressor of NETosis in health not only unveil the underlying mechanism as to why neutrophils are predisposed to NETosis in diabetes, but also identified a natural, endogenous NETosis inhibitor that can be readily (re-)activated as an anti-NETosis treatment. Repurposing natural compounds or drugs to boost SIRT1 activity in the neutrophils can be a new generation of therapeutics to alleviate NET-mediated diabetic complications, and inflammation in general.

Our findings on the inhibitory role of a longevity protein, SIRT1, on diabetes-exacerbated NETosis present a new concept of preventing inflammation from the perspective of “anti-aging”. “Aged” neutrophils are more prone to NETosis,[15–16] similar to diabetes. To tackle the ever-growing “diabetes pandemic” and the aging population, SIRT1 may be a bullet that addresses both issues in one shot to lessen the healthcare burden.

## Methods

### Animals

Animal protocols were reviewed and approved by the Institutional Animal Care and Use Committee of Nanyang Technological University, Singapore (Protocol number: A19051). Unless otherwise indicated, male mice (C57BL/6J background) bred in-house by Nanyang Technological University Animal Research Facility or ordered from InVivos (Singapore) and the Jackson Laboratory (USA) were used in this study. Streptozotocin (STZ)-induced diabetic mice (and their age-matched vehicle-treated controls) were used for the study 10- to 12-weeks post STZ induction. Diabetic *db/db* mice (and their corresponding normoglycemic *db/m+* controls) were used between 10 to 12 weeks of age. *Vav1-Cre Padi4^fl/fl^* mice that lack PAD4 in hematopoietic cells were bred in-house from breeding pairs obtained from Denisa Wagner (Boston, MA, USA).

All mice were fed regular chow *ad libitum* and were maintained in specific pathogen free animal facilities.

### Induction of diabetes in mice using STZ

After acclimatization, 5- to 6- weeks old male C57BL/6J mice were induced to be diabetic via multiple low doses of STZ. Briefly, mice were fasted for 6 hours with no restrictions to water intake, followed by intraperitoneal injection of either vehicle (0.1 M sodium citrate buffer, pH 4) or STZ (50 mg/kg per day) for 5 consecutive days. Non-fasting blood glucose levels were monitored biweekly. For STZ-treated mice, only those with non-fasting blood glucose levels above 16 mmol/L for at least 3 checks were considered diabetic and were used for the study.

### Subject recruitment for human blood samples

All protocols were approved by the Institutional Review Board of Nanyang Technological University, Singapore (IRB-2021-01-037, IRB-2021-02- 042) and Domain Specific Review Board, National Healthcare Group, Singapore (NHG DSRB Ref: 2020/01394). The study was conducted in accordance with the WMA Declaration of Helsinki, and in compliance with the Department of Health and Human Services Belmont Report, and the Human Biomedical Research Act (Singapore). Inclusion criteria for healthy controls were age between 21 to 65 years old, self-declared healthy non-smokers with no personal history of diabetes and not having any recent infection in the past 2 weeks. For subjects with diabetes, the inclusion criteria were individuals with HbA1c equaled to or greater than 8%, non-smokers and not having any recent infection in the past 2 weeks. Exclusion criteria for subject recruitment were individuals more than 65 years of age, on steroid, immunosuppressive or anti-inflammatory medications (such as aspirin, ibuprofen and NSAIDs) in the past 2 weeks, showing signs of active infection (indicated by pyrexia, elevated leukocyte counts and diagnosis of infection) in the past 2 weeks, diagnosis of cancer in the past 5 years, overt heart failure, or hospitalized for any condition in the past 1 month. Written informed consent was obtained from all recruited subjects.

Blood was collected by venipuncture from 14 subjects with type 2 diabetes mellitus (6 males, 8 females; age range 34-65 years old; mean age 52.6 years old) and 13 healthy individuals (3 males, 10 females; age range 35-58 years old; mean age 47.2 years old) (Table S1, Supporting Information). Subjects with diabetes (duration ranged from 1 to 28 years, mean duration 12.93 years, median duration 12.5 years) had co-morbidities such as hypertension and hyperlipidemia and were on anti-diabetic medications such as metformin and statin. Patients with HbA1c equaled to or greater than 8% (reflecting non-compliance or non-responders) were recruited as there were no drug naïve patients available.

### Human neutrophil isolation

Neutrophils were isolated by density gradient centrifugation upon receipt of the fresh blood samples (each about 12 mL) collected in EDTA-coated vacutainers.[7] For each sample, blood was loaded onto Histopaque-1119 (Sigma) and centrifuged at 1,100 xg for 21 minutes (acceleration: 2; brake: 1) at room temperature. The leukocyte layer with significantly reduced amount of red blood cells was transferred to HBSS (without Ca^2+^ and Mg^2+^) for a wash, resuspended again in HBSS and layered on top of a gradient column of Percoll (Cytiva) dilutions at 85%, 80%, 75%, 70% and 65% with 85% at the bottom of the column. The column was then centrifuged at 1,100 xg for 22 minutes (acceleration: 2; brake: 1) at room temperature. Neutrophils were harvested from the milky layer at the 75/80% and 70/75% interfaces, spun down at 400 xg for 10 minutes, and resuspended in a suitable medium for live cell assays or snap frozen for Western blotting. Wright-Giemsa staining confirmed purity of cells was >99%.

### Mouse neutrophil isolation

Each mouse was terminally bled under deep anesthesia for 1 mL into the anti-coagulation buffer (15 mM EDTA and 1% BSA in 1x PBS; sterile filtered) via retro-orbital plexus and neutrophils were isolated using Percoll (Cytiva) gradients.[7] In brief, each mouse blood sample was centrifuged at 500 xg for 12 minutes at room temperature. Supernatant containing the plasma was discarded and blood cells were resuspended in the anti-coagulation buffer for layering onto a gradient column of Percoll (Cytiva) dilutions at 78%, 69%, and 52% with 78% at the bottom of the column. The column was centrifuged at 1,500 xg for 32 minutes (acceleration: 3; brake: 0) at room temperature. Cells at the 78%/69% interface were collected and subjected to a brief red cell lysis in a hypotonic solution. Neutrophils were then spun down and resuspended in a suitable medium for live cell assays. Purity of cells was >95% as reflected by Wright-Giemsa staining.

### NETosis assay

Human and mouse blood neutrophils were resuspended in RPMI-1640 medium supplemented with 10 mM HEPES after isolation and were plated at 1 x 10^4^ cells/well in 96-well CELLBIND plates (Corning). Where appropriate, cells were pre-treated with Ex-527 (10 μM, 30 minutes), RSV (50 μM, 1 hour) or SRT2104 (3 μM, 1 hour) before stimulation for NETosis. For human neutrophils, cells were stimulated with 4 μM ionomycin (ThermoFisher Scientific) for 4 hours (for NETosis quantification) or 5 minutes (for staining citrullinated histone H4, H4Cit, a marker for PAD4 activity). For mouse neutrophils, cells were stimulated with 10 μg/mL LPS (*Klebsiella pneumoniae*, Sigma) for 4 hours (for NETosis quantification) or 2.5 hours (for H4Cit staining). Cells were subsequently fixed in 2% paraformaldehyde (PFA) (Electron Microscopy Sciences) containing Hoechst 33342 (1:10,000, Invitrogen) at 4°C overnight. For evaluation of H4Cit, cells fixed in 2% PFA were permeabilized, incubated with anti-H4Cit (1:500, Millipore, 07- 596) and then with Alexa Fluor 488-conjugated secondary antibody (1:1,500, Invitrogen). Treatments were conducted in duplicates per experiment and ten non-overlapping images of each well were acquired on an Axio Observer 7 (Zeiss) coupled to a Hamamatsu Orca Flash 4.0 sCMOS camera (Hamamatsu) using a LD-Plan Neofluar 20x/0.4 Corr Phase Contrast objective (Zeiss). Exposure intensity and duration were kept constant for all treatments of the same plate. NETs, defined as Hoechst 33342-stained cells with “tails” or streaks (spread NETs) (Figure S1, Supporting Information), were manually counted in black-and-white images. H4Cit^high^ cells were quantified (after immunostaining for H4Cit) by manual thresholding to create a mask using ImageJ / Fiji for automatic cell counting, with exclusion of small particles [Size (inch^2): 0.02-infinity]. The same manual thresholding value was adopted for all images acquired in the same set of experiment.

### Culture and differentiation of HL-60 cells

HL-60 cells (ATCC, CCL-240) were cultured in RPMI- 1640 medium supplemented with 15% fetal bovine serum (Sigma), 1% penicillin/streptomycin (Life Technologies) and 10 mM HEPES (Life Technologies) (complete RPMI). Cells were passaged every 3 days and maintained at a density around 0.8 x 10^6^ cells/mL. To obtain neutrophil-like granulocytes, HL-60 cells were differentiated with 1.3% DMSO (Sigma, D2650) for 6 days and used for subsequent experiments. Routine testing indicated the cell line was free from mycoplasma contamination.

### Adapting HL-60-derived neutrophils (dHL-60 cells) for hyperglycemia study

On day 5 of differentiation, dHL-60 cells were either cultured in 1.3% DMSO-supplemented complete RPMI containing 11 mM glucose (basal glucose, BG) or 44 mM glucose (high glucose, HG) for 24 hours. Complete RPMI supplemented with 1.3% DMSO and mannitol (33 mM mannitol) was included as a control to exclude any effect from osmotic pressure. Cells were then changed into assay medium (RPMI-1640 with 10 mM HEPES), treated with Ex-527 (10 μM, 30 minutes), RSV (50 μM, 1 hour) or SRT2104 (3 μM, 1 hour), where appropriate, before stimulated with 4 μM ionomycin (ThermoFisher Scientific) for 4 hours (for quantification of NETs) or 1.5 hours (for H4Cit) before fixation in 2% PFA containing Hoechst 33342 (1:10,000) at 4°C overnight. Acetylated histone H3 of dHL-60 cells with or without high-glucose exposure was probed with an acetyl-histone H3 antibody (1:500, Sigma-Aldrich, 06-599). Other post-fixation processing, microscopy and image analysis were performed as aforementioned.

### SIRT1 activity assay

Immunoprecipitation for total SIRT1 protein from equal number of cells (1 x 10^6^ cells) was first performed as previously described.[44] Briefly, cells were lysed in M-PER buffer (ThermoFisher Scientific) supplemented with Halt protease and phosphatase inhibitor cocktail (1:100, ThermoFisher Scientific) at 4°C for 1 hour. The supernatant was then incubated for 30 minutes at room temperature with Dynabeads (Life Technologies) pre-conjugated with a SIRT1 antibody (Millipore, Cat. no. 07-131). After washes with 200 μL washing buffer, 5 μL of the eluates was then used for a fluorometric SIRT1 activity assay (Abcam, ab156065). SIRT1 activity was measured at 1-minute interval for 20 minutes for human neutrophils and dHL-60 cells. Linear regression was performed and SIRT1 activity of each sample was determined from the slope of the linear regression analysis.

### Western blot analysis

For whole cell lysates, cells were snap-frozen in liquid nitrogen, followed by homogenization in M-PER buffer (ThermoFisher Scientific) supplemented with Halt protease and phosphatase inhibitor cocktail (1:100, ThermoFisher Scientific) at 4°C for one hour. After centrifugation at 14,000 xg for 15 minutes at 4°C, bicinchoninic acid protein assay (Invitrogen) was performed to measure the protein content of the supernatant. Equal amount of protein was resolved on gradient gels (Bolt 4-12% Bis-Tris Plus gels, Life Technologies) and transferred on to PVDF membranes. For samples immunoprecipitated for SIRT1, bicinchoninic acid protein assay was also conducted to enable equal loading of protein, which was resolved on Bolt 8% Bis-Tris gels (Life Technologies) and transferred on to PVDF membranes. After blocking, the blots were incubated with rabbit polyclonal anti-SIRT1 (Cell Signaling Technology, #9475, for human neutrophils; Abcam, ab189494, for dHL-60 cells) or mouse monoclonal anti-PADI4 (Abcam, ab128086) at 4°C overnight, and subsequently with an appropriate HRP-conjugated secondary antibodies (1:10,000) for 2 hours at room temperature. Blots were developed with enhanced chemiluminescence substrate (Advansta). GAPDH blotting serves as a loading control. Images acquired were analyzed with ImageJ (NIH, Version 1.53).

### SIRT1 knockdown using small interfering RNA (siRNA)

On day 5 of differentiation, dHL-60 cells were subjected to siRNA transfection according to manufacturer’s instructions (Lonza). Briefly, 2 x 10^6^ cells were electroporated with 0.1 nmol scrambled siRNA (Sigma, SIC001) or 0.1 nmol siRNA targeting human SIRT1 (Sigma, SASI_HS01_0015-3666). Cells were then cultured in complete RPMI supplemented with 1.3% DMSO for another 24 hours. Cells were checked for knockdown efficiency using Western blotting (anti-SIRT1, Abcam, ab189494) and examined for NETosis.

### Confocal microscopy for SIRT1 and PAD4 localization

Basal glucose-cultured or high glucose-exposed dHL-60 cells were seeded in the µ-Slide 8-well chambers (ibidi). Upon settled for 15-20 minutes, cells were fixed in 2% PFA (Electron Microscopy Sciences) at room temperature for 10 minutes. Cells were then permeabilized, blocked, and incubated with primary antibodies (anti-SIRT1, Merck, 07-131, 1:1,000; anti-PADI4, Abcam, ab128086, 1:2,000) at 4°C overnight. After washes, cells were incubated with corresponding secondary antibodies conjugated with Alexa Fluor 488 and Alexa Fluor 555 (1:1,500 each, Invitrogen). Single-plane images were acquired on an LSM800 laser scanning confocal microscope (Carl Zeiss) equipped with a Plan Apochromat 63x/1.4 oil objective. Exposure intensity and exposure duration for each channel were kept constant for all cells on the same µ-Slide.

### Super-resolution imaging on chromatin conformation

Human neutrophils (healthy individuals versus diabetic subjects) were seeded on coverslips and exposed to SIRT1 inhibitor (10 μM Ex-527, 30 minutes), SIRT1 activator (50 μM RSV or 3 μM SRT2104, 1 hour) or PAD4 inhibitor (10 μM BB-Cl-amidine, 30 minutes). BB-Cl-amidine was added 30 minutes prior to Ex- 527 incubation when used in combination. Cells were then fixed in 2% paraformaldehyde for 10 min at room temperature, stained with Hoechst 33342 (1:2,000) for 15 min at room temperature and mounted in Fluro-Gel (Electron Microscopy Sciences). Single-plane nuclear images of 20 cells were acquired on a Nikon ECLIPSE Ti2-E AXR confocal microscope coupled to the Nikon Spatial Array Confocal (NSPARC) detector and GaAsP PMT using a CFI Plan Apochromat Lambda D 60x/1.42 oil objective with Nyquist sampling. Acquisition settings were kept constant for all cells. Chromatin compaction states was blindly evaluated by a trained investigator using the NIS-Elements raw files opened on ImageJ (NIH, Version 2.14.0). Chromatin was defined as compact if heterochromatin could be clearly visualized lining the rim of the nucleus (Figure S4, top panels, Supporting Information). Chromatin that appeared more diffused in the nucleus and at the rim was defined as decondensing chromatin with a less compact conformation (Figure S4, bottom panels, Supporting Information).

### Statistical analyses

All data are presented as mean ± standard error of mean (SEM). All experiments were performed for at least two independent runs. Data normality was examined using Shapiro-Wilk test prior to further analyses. Normally distributed data was analyzed using unpaired two-tailed Student’s t-test, one-way ANOVA, or two-tailed Pearson correlation, while Mann-Whitney test or two-tailed Spearman correlation was adopted if otherwise. Statistical analyses were performed using GraphPad Prism (Version 9.4.1). P < 0.05 is considered statistically significant. Sample sizes were determined based on published work and prior knowledge of human studies and animal models.

## Acknowledgements

We thank Mohammad Firdaus Bin Mohd Yusoff and Chung Yin Daryl Yuen for pilot experiments. The study was supported by Nanyang Assistant Professorship (The Lee Kong Chian School of Medicine, Nanyang Technological University Singapore Start-Up Grant), the Singapore Ministry of Education under its Singapore Ministry of Education Academic Research Fund Tier 1 (RG25/20), the Singapore Ministry of Health’s National Medical Research Council under its Open Fund - Individual Research Grant (MOH-001495), and Vascular Research Initiative (022415- 00001, Lee Kong Chian School of Medicine Strategic Academic Initiative) (to S.L.W.). L.D.W. was supported by the Nanyang President’s Graduate Scholarship from Nanyang Technological University Singapore.

## Author contributions

L.D.W and S.L.W. designed and conducted experiments, acquired, analyzed and interpreted data, and wrote the manuscript. R.D. and H.W.H. recruited human subjects and interpreted data. S.L.W. acquired funding for the research and supervised the study. All authors critically reviewed the manuscript and approved the manuscript for submission.

## Data availability

The data that support the findings of this study are available on reasonable request from the corresponding author.

