## Supplemental figures and table for "Sirtuin 1 is an endogenous NETosis inhibitor that becomes dysfunctional in diabetes"

**The Appendix file contains 4 figures and 1 table –**

**Figure S1.** Representative image of human neutrophils stimulated for NETosis.

**Figure S2.** Validation of the dHL-60 cell model for *in vitro* study of the impact of hyperglycemia on NET formation.

**Figure S3.** Treatment of β-nicotinamide adenine dinucleotide (β-NAD, 100 μM, 30-minute pre-incubation) did not influence NET formation.

**Figure S4.** Compact and more decondensed chromatin conformations as observed using super-resolution microscopy.

**Table S1.** Clinical readouts of healthy individuals and subjects with diabetes analyzed using unpaired two-tailed Student's t-test (unless otherwise stated).

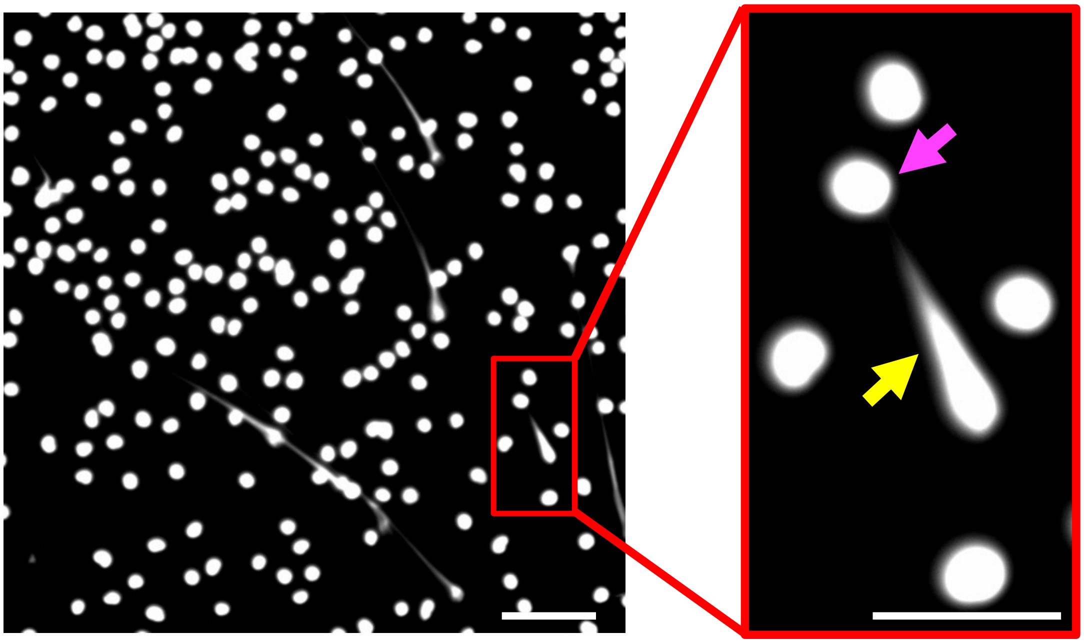

**Figure S1. Representative image of human neutrophils stimulated for NETosis.** (Left) A representative example for NET quantification is presented. Scale, 100 μm. (Right) Zoom-in inset of the image for quantification (red box). Yellow arrow indicates a cell that has undergone NETosis and the magenta arrow indicates a cell without NET release. Scale, 50 μm.

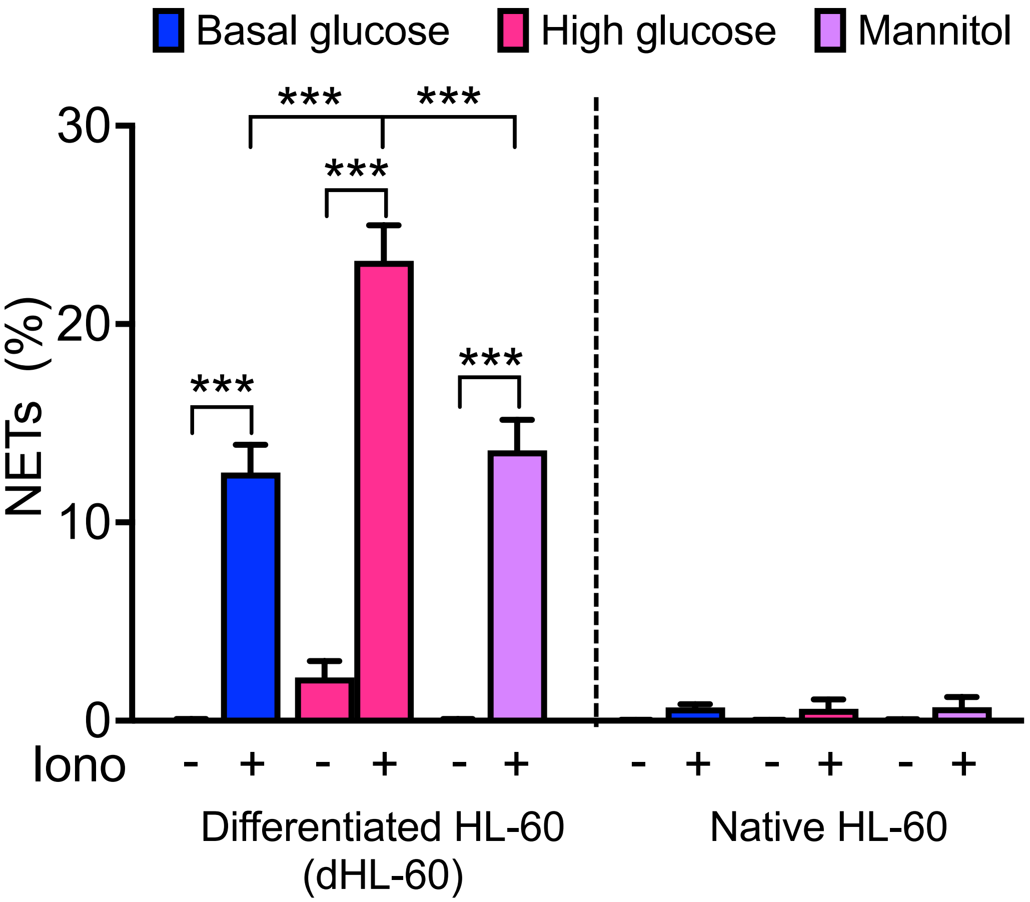


**Figure S2. Validation of the dHL-60 cell model for *in vitro* study of the impact of hyperglycemia on NET formation.** Experiments were performed by two independent investigators with robust, aligning observations. Data were analyzed by Mann-Whitney test, n = 13 biological replicates per group, ***P<0.001. Iono, ionomycin. Data are mean ± SEM.

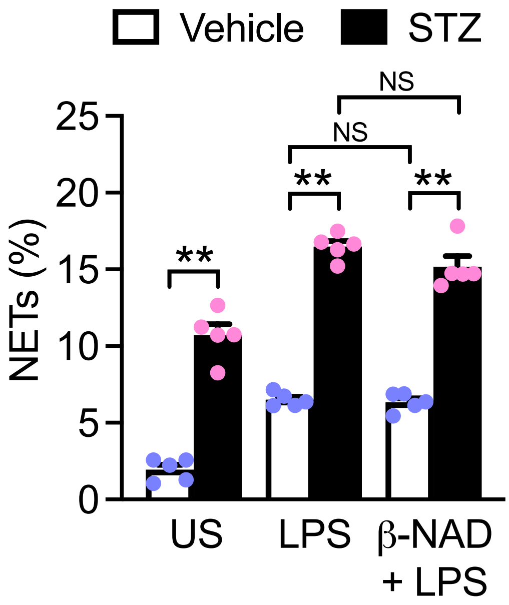


**Figure S3. Treatment of β-nicotinamide adenine dinucleotide (β-NAD, 100 μM, 30-minute pre-incubation) did not influence NET formation.** Data were analyzed by Mann-Whitney test, n = 5 mice per group, **P<0.01, NS, non-significant. US, unstimulated; LPS, lipopolysaccharides. Data are mean ± SEM.

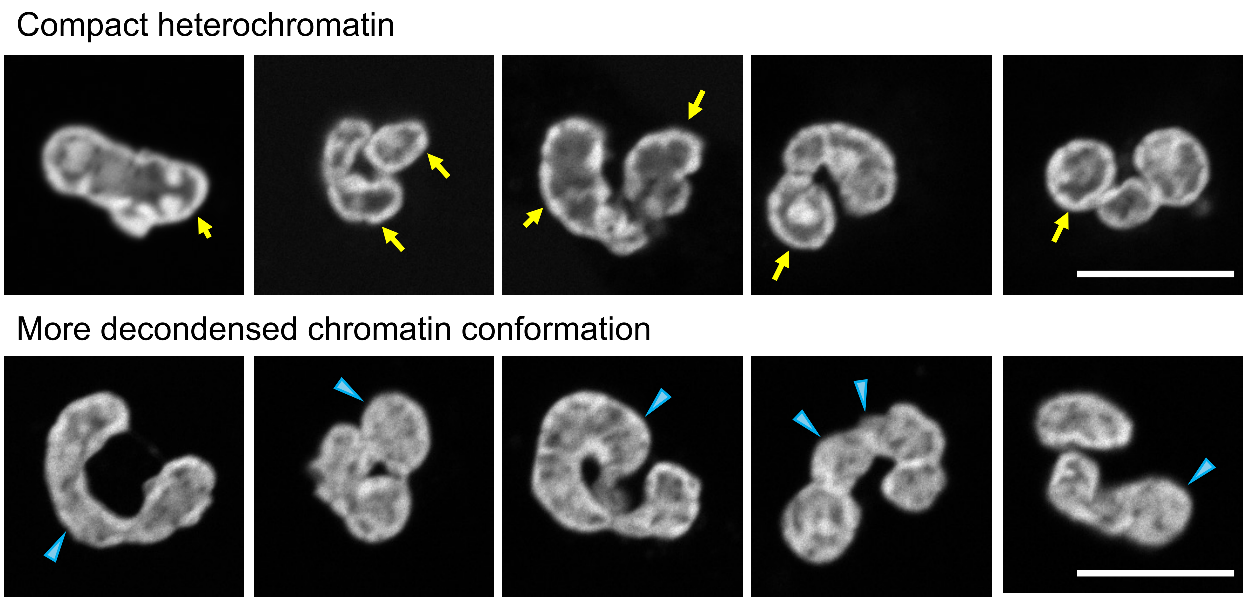


**Figure S4. Compact and more decondensed chromatin conformations as observed using super-resolution microscopy.** Single-plane nuclear images of intact human neutrophils were acquired using super-resolution microscopy and used as a training set for the evaluation of chromatin compaction states of neutrophils by a blinded investigator (Figure 6E). Chromatin was determined to be compact when clear heterochromatin lining the rim of the nucleus (indicated by yellow arrows) could be observed. In comparison, chromatin that appeared more diffused in the nucleus and at the rim (indicated by blue arrowheads) was determined as decondensing chromatin with a less compact conformation. Scale, 10 µm.

**Table S1. Clinical readouts of healthy individuals and subjects with diabetes analyzed using unpaired two-tailed Student's t-test (unless otherwise stated)**

| | Healthy individuals | Diabetic patients | P value |
|---|---|---|---|
| **Age (years)** | 47.15 ± 2.380 | 52.57 ± 2.357 | 0.1188 |
| **Leukocyte count (×$10^3$/µL)^** | 5.600 ± 0.4171 | 7.357 ± 0.4591 | 0.0074 |
| **Neutrophil count (×$10^3$/µL)** | 3.118 ± 0.2735 | 4.474 ± 0.3757 | 0.0080 |
| **Lymphocyte count (×$10^3$/µL)** | 1.779 ± 0.1983 | 2.114 ± 0.1492 | 0.1859 |
| **Platelet count (×$10^3$/µL)^** | 261.4 ± 17.28 | 257.8 ± 22.19 | 0.9708 |
| **HbA1c (%)** | 5.531 ± 0.1247 | 9.743 ± 0.4826 | <0.0001 |
| **Fasting glucose (mmol/L)** | 5.092 ± 0.1332 | 12.15 ± 1.070 | <0.0001 |
| **Total cholesterol (mmol/L)** | 5.277 ± 0.2918 | 4.479 ± 0.2141 | 0.0351 |
| **Triglycerides (mmol/L)^** | 1.515 ± 0.5809 | 2.171 ± 0.3615 | 0.0109 |
| **HDL-C (mmol/L)** | 1.508 ± 0.1380 | 1.329 ± 0.07944 | 0.2627 |
| **LDL-C (mmol/L)** | 3.246 ± 0.2371 | 2.400 ± 0.2029 | 0.0122 |

Data normality was determined with Shapiro-Wilk test and data are presented as mean ± SEM. Samples from the recruited subjects (13 healthy and 14 diabetic) were analyzed.
^Mann-Whitney test was adopted for analyses of leukocyte count, platelet count, and triglycerides as data was not normally distributed.
